# Soil salinity limits plant shade-avoidance

**DOI:** 10.1101/289124

**Authors:** Scott Hayes, Chrysoula K. Pantazopoulou, Kasper van Gelderen, Emilie Reinen, Adrian Louis Tween, Ashutosh Sharma, Michel de Vries, Salomé Prat, Robert. C. Schuurink, Christa Testerink, Ronald Pierik

## Abstract

Global food production is set to keep increasing despite a predicted decrease in total arable land [1]. To achieve higher production, denser planting will be required on increasingly degraded soils. When grown in dense stands, crops elongate and raise their leaves in an effort to reach sunlight, a process termed shade-avoidance [2]. Shade is perceived by a reduction in the ratio of red (R) to (FR) light and results in the stabilisation of a class of transcription factors known as PHYTOCHROME INTERACTING FACTORs (PIFs) [3,4]. PIFs activate the expression of auxin biosynthesis genes [4,5] and enhance auxin sensitivity [6], which promotes cell wall loosening and drives elongation growth. Despite our molecular understanding of shade-induced growth, little is known about how this developmental programme is integrated with other environmental factors.

Here we demonstrate that low levels of NaCl in soil strongly impair the ability of plants to respond to shade. This block is dependent upon abscisic acid (ABA) signalling and the canonical ABA signalling pathway. Low R:FR light enhances brassinosteroid (BR) signalling through *BRASSINOSTEROID SIGNALLING KINASE 5* (*BSK5*), and leads to the activation of BRI1 EMS SUPPRESSOR 1 (BES1). ABA inhibits *BSK5* up-regulation and interferes with GSK3-like kinase inactivation by the BR pathway, thus leading to a suppression of BES1:PIF function. By demonstrating a link between light, ABA and BR-signalling pathways this study provides an important step forward in our understanding of how multiple environmental cues are integrated into plant development.

## Results and Discussion

### Soil salinity inhibits plant shade-avoidance

Soil salinity is detrimental to plants and often results in a reduction in stem and root biomass accumulation [7]. Most previous studies have, however, focused on the effects of salt at a range of very high concentrations (at a median concentration of 150mM NaCl) [8]. We opted for a nuanced approach to investigate how low concentrations of NaCl may affect plant shade-avoidance. Growing plants in tissue culture often masks genes involved in NaCl-sensitivity [9,10], and so for phenotypic experiments plants were grown on soil. Arabidopsis thaliana (arabidopsis) seeds were germinated under white light (Wl) for three days, before transfer to new soil that had been pre-treated with NaCl solution. Following this, plants were returned to Wl for a further day to acclimate, before they were shifted to Wl or Wl with supplementary far-red LEDs (+FR). Hypocotyl lengths were measured on day seven (for a schematic diagram of the treatments see Figure S1A). Water or NaCl solution was applied from below and the soil was kept saturated to avoid dehydration.

We observed a strong NaCl-mediated reduction in +FR-induced hypocotyl elongation when plants were watered with as low as 10mM NaCl (Figure 1A). Increasing NaCl concentrations beyond 25mM to 75mM NaCl provided no further inhibition of +FR-induced hypocotyl elongation and so 25 mM NaCl was selected for further investigation. NaCl inhibited +FR-induced elongation across the 3 days of +FR treatment (Figure S1B). In adult plants, 25 mM NaCl exerted a strong inhibition of +FR-induced rosette expansion, mostly through an inhibition of lamina elongation (Figures 1B, 1C, S1C and S1D). Additionally, we found that NaCl inhibited +FR induced elongation in tomato and tobacco seedlings (Figure 1D and 1E), raising the possibility that this is a general phenomenon in shade-avoiding plants.

**Figure 1.**
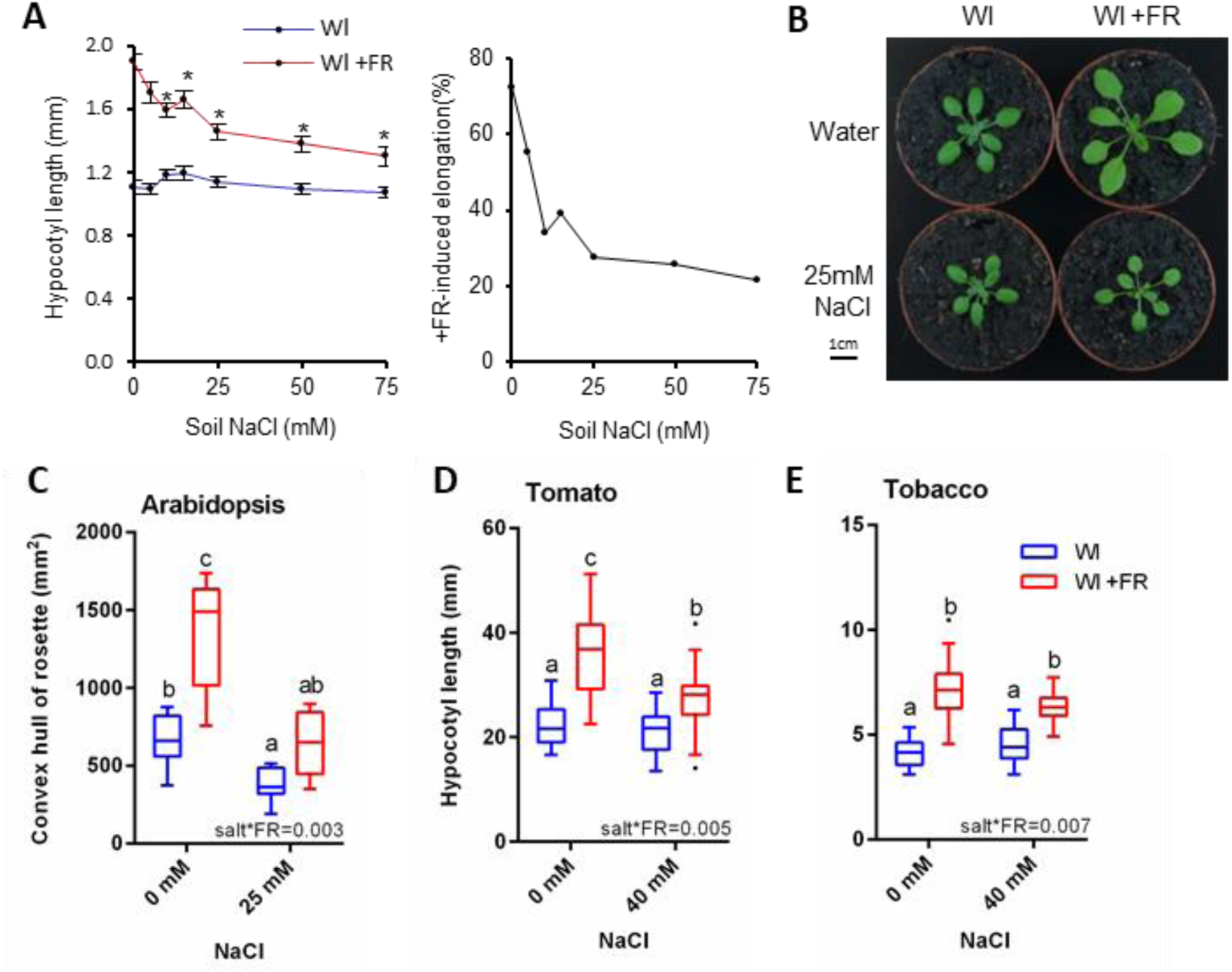
Soil salinity inhibits +FR-induced elongation (See also Figure SI) **(A)** Left: Hypocotyl length of 7-day old Col seedlings germinated in Wl, transferred to NaCI soil of the indicated concentration at day 3 and then shifted to WL ±FR at day 4. Data represent mean (n>19) ± SE. Asterisks indicate salt-mediated inhibition of hypocotyl elongation compared to OmM NaCI control-Students t-test (p < 0.05). Right: The same data expressed as the % +FR-induced elongation at each salt concentration. **(B)** Representative 21-day old Col plants germinated in Wl and transferred to ±25mM NaCI at day 3, before shifting to WL ±FR at day 10. **(C)** Rosette circumference of plants grown as in “B” (n>10). **(D)** Hypocotyl length of 11 day old tomato *(Solanum lycopersicum* var. “Moneymaker”) seedlings germinated in Wl, transferred to ±40mM NaCI at day 7, and shifted into WL ±FR at day 8 (n>19). **(E)** Hypocotyl length of tobacco *(Nicotiana benthamiana*) seedlings grown as in “D” (n>18). Box plots are visualised by the Tukey method. In all figures, different letters designate significantly different means by 2-way ANOVA + Tukey’s post-hoc test (p <0.05). Interaction P value is shown inset.

### Soil salinity acts through the ABA pathway

Many plant responses to soil salinity are mediated through the hormone ABA [11]. We found that transcripts of *SAG29* (an ABA-responsive gene [12]) accumulated to high levels in +NaCl +FR conditions (Figure 2A). This accumulation did not occur in plants lacking four ABA-responsive transcription factors (*ABA-responsive element-binding factor 1, 2, 3* and *4* [13], here referred to as *arebQ*). We therefore decided to assess whether ABA signalling is required for NaCl-mediated inhibition of +FR-induced elongation. In mutants that lacked either four of the ABA receptors (*pyr1*/*pyl1*/*pyl2*/*pyl4*-here referred to as *abaQ* [14]) or two ABA-signalling kinases (*Sucrose non-fermenting 1-Related protein Kinase 2.2* and *2.3*-*snrk2.2snrk2.3*) [15], NaCl no longer had an effect on +FR-induced hypocotyl elongation (Figure 2B-C). Whereas plants lacking a single ABA activated transcription factor, *ABA INSENSITIVE 5* (*abi5-1*) still showed a strong NaCl-mediated inhibition of hypocotyl elongation under +FR light (Figure S2A), plants lacking the *AREB* quartet (*arebQ*) elongated to the same extent in the presence and absence of NaCl (Figure 2D). Taken together, these results suggest that the effect of NaCl on +FR-induced elongation is dependent upon enhanced ABA synthesis or signalling.

**Figure 2.**
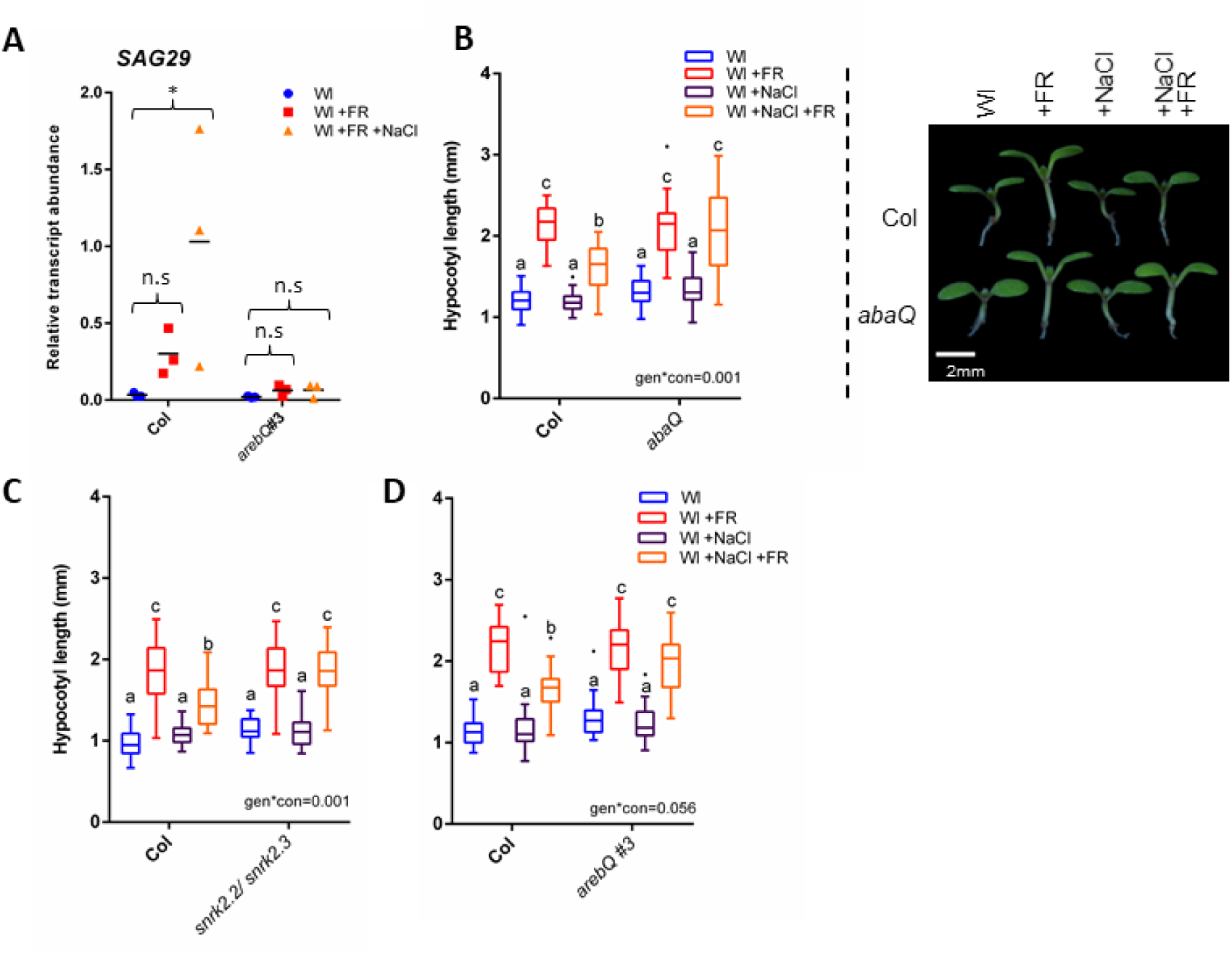
Salinity-mediated inhibition of +FR-induced elongation requires ABA signalling (See also Figure S2) **(A)** Relative *SAG29* transcript abundance in the hypocotyls of Col and *arebQ* mutant plants grown 3 days in Wl, transferred to ±25mM NaCI soil at day 3 and then shifted to WL ±FR at day 4. Tissues were harvested at Zt 4.5 on day 6 (n=3). *= significant difference from Wl control. Hypocotyl lengths of 7 day old wild type and **(B)** *abaQ,* **(C)** *snrk2.2/snrk2.3* and **(D)** *arebQ* mutants germinated in Wl, transferred to±25mM NaCI soil at day 3 and then shifted to WL±FR at day 4 (n>22). Right panel of “A” depicts representative seedlings grown in these conditions. Box plots are visualised by the Tukey method. Gene expression studies show individual values, with a horizontal bar representing the mean. Different letters designate significantly different means by 2-way ANOVA+ Tukey’s post-hoc test (p<0.05). Interaction Pvalue is shown inset.

In addition to these mutant and gene expression analyses, we found that applying increasing concentrations of ABA directly between the cotyledons had a similar effect as applying increasing NaCl concentrations to the soil (Figure 1A versus S2B). Low concentrations of ABA provided a strong break on hypocotyl elongation in +FR light, this inhibition was saturated by 10 µM ABA and remained constant until at least 100 µM ABA. As with NaCl, the inhibition of +FR-induced hypocotyl elongation by exogenous ABA was absent in the *abaQ* (Figure S2C) and *snrk2.2*/*snrk2.3* mutants (Figure S2D). Despite the clear requirement for ABA signalling, NaCl still inhibited +FR-induced elongation in mutants that have reduced levels of ABA synthesis, such as *aba2-1* and *aba3-1* (Figure S2E and S2F) and we were unable to detect any increase in ABA levels in whole NaCl-grown seedlings (Figure S2G). It may be that ABA signalling is enhanced only locally, ABA distribution is altered or that there is direct activation of the ABA signal pathway by an unknown mechanism.

### Salt and ABA impede upon PIF4/PIF5 action

We next investigated the manner in which ABA and NaCl inhibit +FR-induced elongation. The induction of hypocotyl elongation by +FR requires PIF4, PIF5 and PIF7 [3,4,16] (Figure 3A). In our conditions PIF7 was the dominant PIF driving +FR-induced hypocotyl elongation, as the *pif7* single mutant showed no significant +FR-induced elongation and there were no additional effects in the *pif4*/*pif5*/*pif7* triple mutant. Single *pif4* or *pif5* mutants appeared similar to the wild type, with strong +FR-induced hypocotyl elongation that was inhibited by NaCl. The *pif4*/*pif5* double mutant did however show reduced +FR-induced elongation, suggesting that these genes act redundantly. Notably, the hypocotyl length of the *pif4*/*pif5* mutant in +FR light was the same as NaCl +FR treated wild type plants and NaCl did not further suppress elongation in this mutant (Figure 3A). In a similar fashion to NaCl, ABA inhibited +FR-induced hypocotyl elongation in the wild type to the same length as the *pif4*/*pif5* mutant, and ABA had no further effect on elongation in this mutant (Figure S2C). These findings suggest that NaCl and ABA could act to suppress PIF4/PIF5 function.

**Figure 3.**
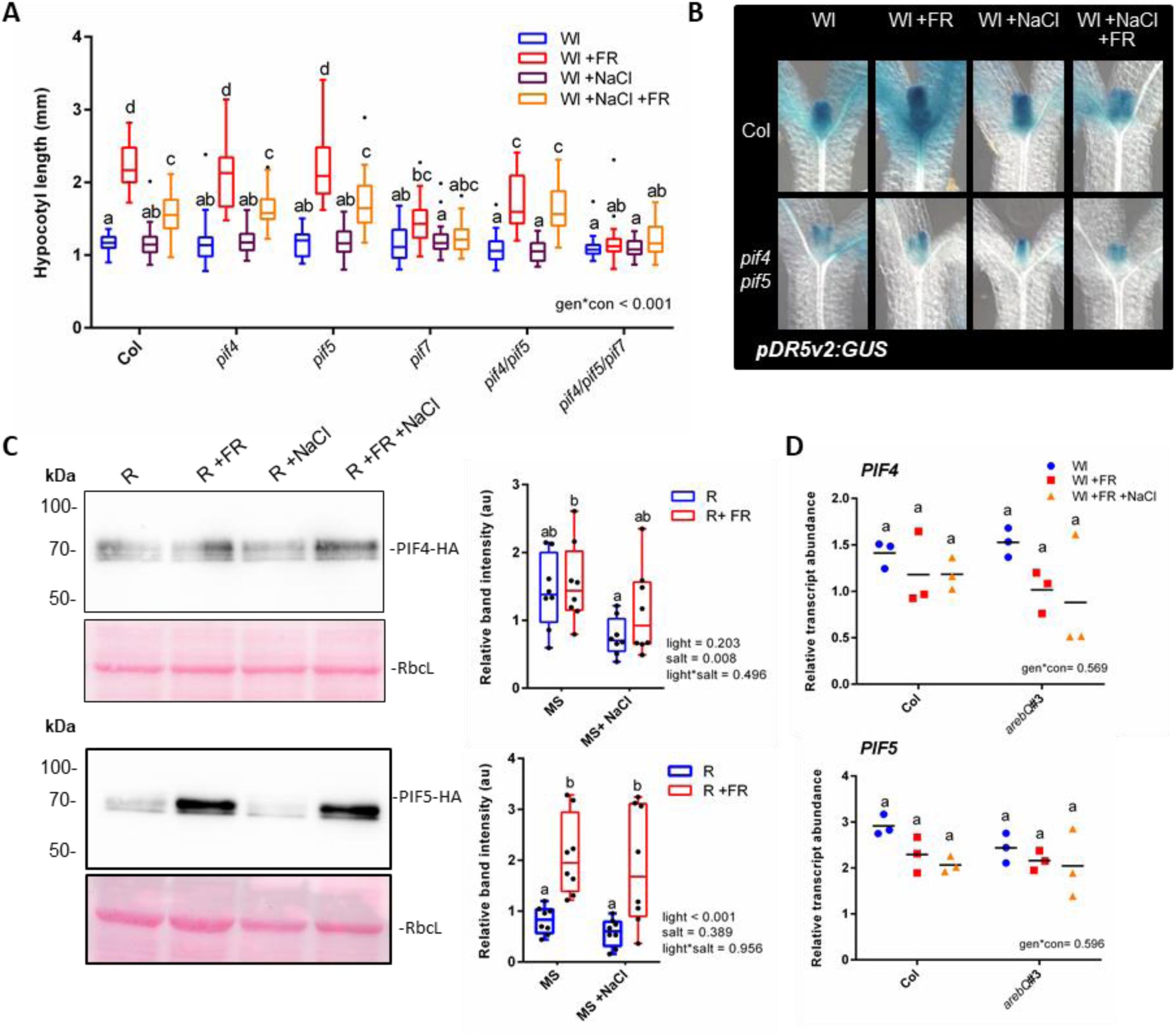
Soil salinity supresses PIF-function and +FR-induced gene expression and auxin signalling (See also Figure S3) **(A)** Hypocotyl length of 7-day old Col, *pif4-101, pif5* and *pif7-l* double and triple mutant seedlings germinated in Wl, transferred to 25mM NaCI soil at day 3 and then shifted to WL ±FR at day 4 (n>23). **(B)** Seedlings expressing a *pDR5v2:GUS* reporter in the Col or *pif4-101/pif5* background were germinated in Wl and transferred to ± 25mM NaCI soil on day 3. On day 4 at plants were shifted to WL ±FR and seedlings were fixed on day 5 at Zt 4.5. Representative hypocotyls of GUS stained seedlings are shown (quantification of the top half of the hypocotyls is shown in Figure 3SE). **(C)** Western blots of *35S:PIF4-HA* (above) and *35S:PIF5-HA* (below) plants germinated on plates in Wl, transferred to 75mM NaCI plates and grown for a further 2 days in R light. At zt 3 on day 5, plants were moved to R ±FR for 1 hour, and seedlings harvested into liquid N_2_. Western blots probed with anti-HA. Pink lanes show ponceau staining of rubisco large subunit (RbcL) as a loading control. Right panel depicts relative band intensity after normalisation to RbcL (n= 8, across 4 separate experiments). **(D)** Relative *PIF4* (above) and *PIF5* (below) transcript abundance in the hypocotyls of Col and *arebQ* mutant plants grown as in “A” that were harvested at Zt 4.5 on day 6 (n=3). Box plots are visualised by the Tukey method (if n >10) or as max/min with every value shown (if n < 10). Gene expression studies show individual values, with a horizontal bar representing the mean. In all figures, different letters designate significantly different means by 2-way ANOVA+Tukey’s post-hoc test (p <0.05).

We found that the expression of two +FR induced genes (*PRE1* and *SAUR16*) was suppressed by pre-treatment with NaCl. In the *pif4*/*pif5* mutant however, NaCl did not affect the expression of these genes (Fig 3SA). We used plants expressing luciferase (LUC) reporter constructs to investigate salt-mediated inhibition of +FR-induced genes over a longer time scale. When LUC expression was driven by the *PIL1* [17] or *IAA19* promoter, +FR light resulted in a rapid induction of luciferase activity and NaCl and ABA caused a sustained suppression of this upregulation over several days (Figure S3B). As both the *PIL1* and *IAA19* promoters contain several G-boxes (Figure S3C) that are established PIF4 and PIF5 direct targets [5,18], these results are consistent with NaCl and ABA-mediated inhibition of PIF4 and PIF5 activity.

PIFs promote hypocotyl elongation in +FR light through an increase in auxin signalling [2]. To investigate whether NaCl and ABA suppressed auxin signalling we generated a DR5v2:GUS auxin reporter line. DR5v2 is a synthetic promoter that was developed as a more sensitive auxin reporter than the widely used DR5 [19]. Three independent DR5v2:GUS lines were tested and confirmed to be highly auxin responsive (Figure S3D). We found that when grown under supplementary FR light, DR5v2:GUS plants had increased GUS staining at the upper part of the hypocotyl, but that this response was attenuated in the presence of NaCl (Figure 3B, S3E). We then crossed this line into the pif4/pif5 mutant background. In the absence of PIF4 and PIF5 we no longer detected a +FR-mediated increase in GUS staining and salt had no further effect on this mutant (Figure 3B, S3E). Again, this is consistent with an NaCl-mediated inhibition of PIF4/ PIF5 activity.

In wild type plants, auxin application promoted hypocotyl elongation in all of our growth conditions, but NaCl still significantly inhibited +FR-induced elongation in the presence of supplementary auxin (Figure S3F), suggesting that NaCl may act to suppress auxin sensitivity or transport [20]. Supplementary auxin also enhanced hypocotyl elongation in the *pif4*/*pif5* mutant in most of the conditions tested, but did not rescue elongation in this mutant to the same length as wild-type plants treated with auxin and +FR (Figure S3F). NaCl had no effect on +FR-induced elongation in the *pif4*/*pif5* mutant, even in the presence of supplementary auxin, which is in accordance with the role of PIF4 and PIF5 in the control of auxin signalling [6].

PIFs are often regulated by light at the level of protein stability [3,21], so we assessed the abundance of constitutively expressed PIF4-HA and PIF5-HA under +FR and +NaCl conditions. While 2-way ANOVAs revealed an effect of NaCl on PIF4 stability, we did not see a statistically significant reduction in PIF4 protein levels in any of the individual light conditions tested (Figure 3C). PIF5 was stabilised by +FR light, but NaCl had no significant effect on this stabilisation (Figure 3C). Additionally, we observed no changes in *PIF4* or *PIF5* expression in NaCl or in the *arebQ* mutant (Figure 3D).

### Brassinosteroid pathway is inhibited by salt and ABA

We hypothesised that PIF4 and PIF5 action could be inhibited through a change in activity of one of their interaction partners. PIF action is suppressed by competitive heterodimerisation with DELLA proteins [22,23]. It has previously been shown that NaCl stabilises DELLA proteins [24] and DELLA degradation is required for +FR-induced shade avoidance to occur [25]. Mutants lacking all five DELLA proteins showed longer hypocotyls than the wild type in all of our conditions (Figure S4A), but still exhibited a strong NaCl-mediated inhibition of +FR-induced elongation. This suggests that NaCl inhibits shade avoidance through another mechanism. A recent study found that the stabilisation of DELLAs by NaCl in the root is transient [26] and so it may be that their role is restricted to growth arrest upon initial NaCl exposure.

In NaCl-treated roots, ABA production precedes a reduction of BR signalling [26] and it is known that BR signalling is necessary for +FR-induced hypocotyl elongation to occur [16]. This raises the possibility that during NaCl exposure ABA acts to inhibit BR signals. We tested the effect of brassinazole (BRZ-an inhibitor of BR synthesis) on hypocotyl elongation under +FR light. We found that BRZ, much like both NaCl and ABA, inhibited hypocotyl elongation very readily at low concentrations and that this inhibition remained constant as concentrations increased (Figure S4B). Intriguingly, hypocotyl length at this plateau was equal to that of +FR-treated plants grown on saline soil in the absence of BRZ. Furthermore, simultaneous application of ABA, BRZ and NaCl had no further effect on the hypocotyl length of +FR-treated plants than NaCl treatment alone (Figure S4C), consistent with these compounds converging on the same pathway. Supporting this, application of epi-brassinolide (BL) rescued +FR-induced hypocotyl elongation in NaCl-exposed plants (Figure S4D). Hypocotyl length was rescued at low concentrations of epi-BL, up to the extent of +FR-treated control plants grown in the absence of NaCl (Figure S4D). We also observed that epi-BL could not rescue hypocotyl elongation in the *yucca2589* mutant (Figure S4E) which probably indicates that in our conditions, BR-mediated rescue of hypocotyl elongation is auxin dependent.

Although it is known that inhibition of BR signalling inhibits shade-avoidance, it is currently unclear whether +FR enhances BR signals. We assessed BR activity through looking at BES1 phosphorylation levels in a *35S:BES1-GFP* line. We found that short term +FR treatments rapidly resulted in a faster running form of BES1 (Figure 4A), which has previously been shown to correspond to the active, de-phosphorylated form of the protein [27]. We found that NaCl inhibited the +FR-mediated induction of this faster running band (Figure 4A), suggesting that BES1 activation by +FR light is inhibited by NaCl. Despite this, we found that plants that hyper-accumulate BES1 (*bes1-D* mutants) actually had shorter hypocotyls under +FR than wild type plants (Figure S4F). Recently it has been shown that the balance between BES1 and PIF4 levels dictates whether BES1 acts as an activator or repressor of BR synthesis genes [28]. Over-accumulation of BES1 suppresses BR synthesis genes and since PIF4 activity is directly suppressed in the absence of BR [29], this may explain the reduced +FR-response in the *bes1-D* line.

**Figure 4.**
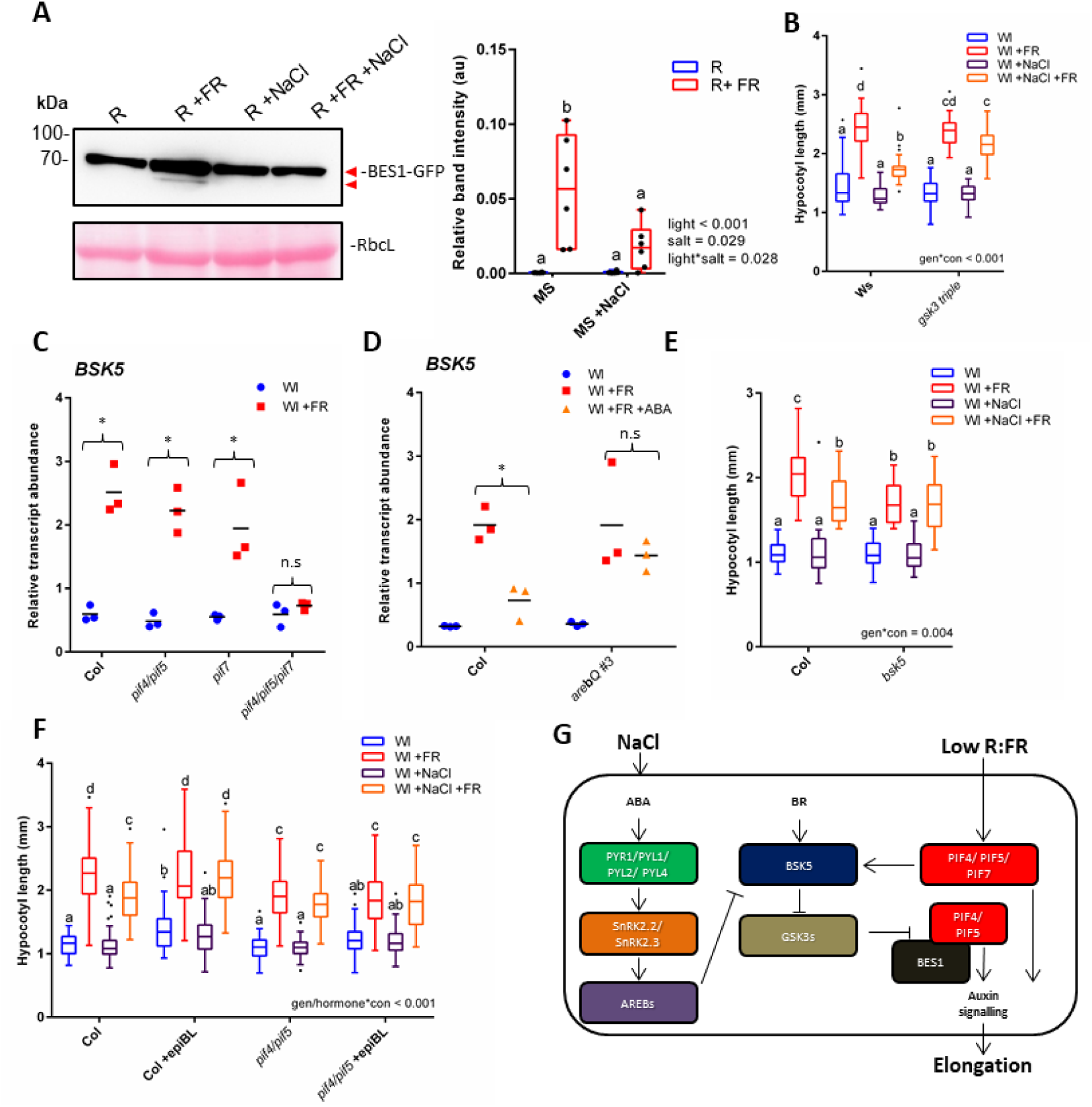
ABA and NaCI inhibit +FR elongation through inhibiting BR signalling (See also Figure S4) **(A)** *35S:BESl-GFP-ex*pressing plants were germinated on plates for 3 days in Wl, transferred to ±75mM NaCI plates and then grown in R light for 2 further days. On day 5, plants were shifted to R ±FR at zt 3 and tissue harvested at zt4. Proteins were identified through Western blots with anti-GFP. Right panel shows quantified lower band intensity relative to RbcL (n=6). **(B)** Hypocotyl length of 7 day-old Ws and *gsk3-triple* mutant germinated in Wl, transferred to ±25mM NaCI soil at day 3 and then shifted to WL±FR at day 4 (n > 24). (C) Col and *pif4-101/pif5, pif7-l* and *pif4-101/pif5/pif7-l* plants were germinated in Wl for 3 days before transfer to new pots. At zt 2.5 on day 7, plants were shifted to Wl ±FR light. At ZT 4.5, the hypocotyls of approximately 20 seedlings per sample were dissected and RNA extracted. *BSK5* transcripts relative to *AT1613320* are shown (n=3). *= +FR-induced increase. **(D)** Col and *arebQ #3* plants were grown as in “C”. At ZT 1.5 on day 7, plants were sprayed with 25 |iM ABA and exposed to Wl ±FR at zt 2.5. At ZT 4.5, the hypocotyls of approximately 20 seedlings per sample were dissected and RNA extracted. *BSK5* transcripts relative to *AT1613320* are shown (n=3). * = ABA-mediated decrease. **(E)** Hypocotyl length of Col and *bsk5* grown as in “B” (n > 22). **(F)** Hypocotyl length of 7 day old Col and *pif4-101/pif5* plants grown as in “B”, with applications 10 |iM epi-BL or ethanol control on days 3-6 (n > 46). **(G)** A proposed mechanism for salt-mediated inhibition of low R:FR-induced elongation. Low R:FR light promotes the stability of PIF4, 5 and 7. These PIFs redundantly upregulate the expression of *BSK5.* BSK5 acts to supress the activity of GSK3-like kinases, which relieves their suppression of the PIF:BES1 signalling module in a positive feedback loop. NaCI promotes the canonical ABA signal transduction pathway, resulting in the increased activity of AREB transcription factors. AREBs inhibit the upregulation of *BSK5* and thereby enhance GSK3-like kinase action. GSK3-like kinases are then free to inhibit the PIF:BES1 signalling module, limiting auxin signalling in the hypocotyl. Box plots are visualised by the Tukey method (if n >10) or as max/min with every value shown (if n < 10). Gene expression studies show individual values, with a horizontal bar representing the mean. Where letters are shown, different letters designate significantly different means by 2-way ANOVA+Tukey’s post-hoc test (p <0.05). Box plots are visualised by the Tukey method. Luciferase data are presented as the median value for each treatment. In all figures, different letters designate significantly different means by 2-way ANOVA + Tukey’s post-hoc test (p <0.05).

BES1 phosphorylation is mediated through BRASSINOSTEROID INSENSITIVE 2 (BIN2-a GSK3-like kinase) which acts as a negative regulator of the BR pathway. Previously it was shown that ABA enhances the kinase activity of BIN2 towards BES1 [30]. We found that mutants that lack BIN2 and two closely related GSK3 kinases (gsk3 triple) no longer showed inhibition of +FR-induced elongation in response to NaCl, ABA or BRZ (Figure 4B, S4G).

### ABA inhibits +FR-induced expression of BSK5

We reasoned that the inhibition of GSK3 kinase function may be required for full +FR-induced hypocotyl elongation. Upstream of GSK3 kinases in the BR signal cascade are the BRASSINOSTEROID SIGNALLING KINASES (BSKs). BSK activation by BR [31] results in the inactivation of GSK3 kinases [31–33]. Despite high redundancy between the 12 BSK family members, [33] mutation of just one (BSK5) alters plant sensitivity to NaCl and ABA [34]. Interestingly, *BSK5* transcripts are enhanced in hypocotyls under +FR light [35]. We found that this upregulation was redundantly regulated by *PIF4, PIF5* and *PIF7* (Figure 4C).

The BSK5 promoter contains a G-box approximately 3.5kb from its start codon (Figure S4H). Multiple PIFs bind directly to this G-box [18], as does at least one AREB family member, ABF3 (in an ABA-dependent manner) [12]. We found that ABA pre-treatment strongly inhibited the +FR-mediated upregulation of *BSK5* transcripts in hypocotyls (Figure 4D) and that ABA-mediated repression of *BSK5* transcripts was dependent upon AREB family transcription factors. +FR-induced hypocotyl elongation was dependent upon *BSK5* as *bsk5* mutants showed only limited +FR-induced elongation (Figure 4E). This elongation was to a similar length as wild-type plants treated with +FR and NaCl and NaCl had no further effects on +FR-induced elongation in a *bsk5* mutant (Figure 4E). These results demonstrate that the +FR-mediated upregulation of *BSK5* is required for maximal hypocotyl elongation under +FR light and they suggest that ABA may at least in-part inhibit +FR-induced elongation through a suppression of *BSK5* transcription.

### Repression of +FR-induced elongation centres on the PIF/BES1 module

Importantly, the rescue of hypocotyl elongation by epi-BL did not occur in the *pif4*/*pif5* mutant, demonstrating that in this context, BR-induced growth requires *PIF4* and/or *PIF5* (Figure 4F). Recent reports have demonstrated a close regulatory relationship between PIF and brassinosteroid signalling. Brassinosteroid upregulates each of the shade-induced genes that we tested (*PIL1, IAA19, SAUR16* and *PRE1*) [29,36,37]. PIF4 heterodimerises with BES1 [28] and a BES1 homologue, BRASINAZOLE RESISTANT 1 (BZR1), and inter-dependently they control the expression of thousands of genes [38]. Additionally, BIN2 has a direct inhibitory effect on both PIF4 [29] and PIF3 [39].

From our results we propose a mechanism (Figure 4G) whereby +FR light promotes the activity of PIF4, PIF5 and PIF7 and these transcription factors then redundantly promote the expression of *BSK5*. BSK5 suppresses GSK3-like kinases, allowing for the activation of the BES1: PIF4/PIF5 signalling module. In saline soils, activation of the ABA signal transduction pathway results in the enhanced action of AREB/ ABF transcription factors that suppress *BSK5* transcription, and therefore break the +FR-induced PIF/BSK5/PIF feed-forward loop. The absence of BSK5 would allow for strong GSK3-like kinase action, leading to a suppression of the BES1: PIF4/PIF5 signal module. We do not rule out other mechanisms by which NaCl may inhibit +FR-induced elongation; indeed we found that mutants lacking the evening complex component *ELF3* had elongated hypocotyls in all of our growth conditions and appeared to lack NaCl-mediated inhibition of +FR-induced hypocotyl elongation (Figure S4H). This implies that NaCl may also supress PIF4 and PIF5 through the circadian clock.

There may be an adaptive significance to our finding that plants grown in saline soils suppress BES1: PIF signalling when they are presented with shade cues. Indeed, a recent study suggested that overexpression of *PIF4* may reduce plant survival in saline conditions [40]. It is possible then that shade avoidance signalling is detrimental to salt survival. It is notable that BR-based chemicals are currently being developed as treatments for salt and drought afflicted crops [41]. It may be that under concurrent salt and shade conditions, re-activation of the BR signal cascade is damaging to plant health.

## Supporting information

Key resources table

Oligonucleotides used in this study

## Author Contributions

Conceptualization, S.H, C.T and R.P; Methodology, S.H, A.T, S.P and R.P; Formal analysis, S.H; Investigation, S.H, C.K.P, K.vG, A.T, E.R, A.S, M.dV and S.P; Resources, R.S, S.P and R.P; Writing (original), S.H; Writing (review and editing), S.H, C.K.P, K.vG, C.T, S.P, R.S and R.P; Visualisation, S.H; Funding acquisition, S.H, R.P and S.P.

## Acknowledgements

We would like to thank Prof. Dolf Weijers (WUR, Wageningen, NL) for providing the *pUC57:DR5v2* plasmid, Prof. Christian Fankhauser (UNIL, Lausanne, CH) for the *pCF402* and *pCF404* plasmids, Prof. Kazuko Yamaguchi-Shinozaki (The University of Tokyo, Tokyo, Japan) for *arebQ* seeds, Prof. Keara Franklin (The University of Bristol) for the *p35S:PIF4-HA* and *p35S:PIF5*-HA seeds ahead of publication, the Plant Ecophysiology Group (Utrecht University, Utrecht, NL) for help with harvesting and gene expression studies and Dr. Charlotte Gommers (WUR, Wageningen, NL) for critical reading of this manuscript. S.H was supported through a fellowship from the European Molecular Biology Organisation (EMBO-ATFL 404-2015) and Marie Sklodowska Curie Actions (792624). The research was funded through the Netherlands Organisation for Scientific Research (NWO-awarded to R.P., VIDI-864.12.003), the Spanish Ministry of Economy and Competitiveness (MINECO-awarded to S.P., BIO2017-90056-R) and the UK Biotechnology and Biological Sciences Research Council (BBSRC, grant BB/M008711/1 to support A.S).

## Methods

### Contact for Reagent and Resource Sharing

Further information and requests for resources and reagents should be directed to and will be fulfilled by the Lead Contact, Ronald Pierik. Correspondences should be addressed to.

### Experimental Model and Subject Details

The main experimental organism used in this study was *Arabidopsis thaliana*. Several mutant arabidopsis lines were used: *pif4-101, pif5* (*pil6-1*), *pif7-1, pif4-101*/*pif5* (*pil6-1*), *pif4-101*/*pif5* (*pil6-1*)/ *pif7-1, abaQ* (*pyr1-1*/*pyl1-1*/*pyl2-1*/*pyl4-1*), *snrk2.2*/*snrk2.3, arebQ* (*areb1*/ *areb2*/ *abf3*/ *abf1-1*), *bsk5, aba2-1, aba3-1, pPIL1:LUC* and *p35S:BES1-GFP* in the Columbia ecotype; *abi5-1* and *gsk3-triple* (*bin2-3/bil1-1/bil2-1*) in the Ws ecotype; and *della global* (*gai-t6*/ *rga-t2*/ *rgl1-1*/ *rgl2-1*/ *rgl3-4*), *p35S:PIF4-HA* and *p35S:PIF5-HA* in the L*er* ecotype. The other species used in this study were tomato (*S.lycopersicum* var. “Moneymaker”) and tobacco (*N. benthamiana*). See the Key Resources Table for details.

### Method Details

#### Morphological studies

Seeds were sown directly onto wetted soil and stratified at 4°C in darkness. After 3-4 days of stratification, plants were moved to growth chambers with a 16:8 hour photoperiod (PP), 130-140 µmol m^-2^ s^-1^ white light. After 3 days, plants were transferred to new soil that had been previously wetted with deionised water or the indicated concentration of NaCl and then returned to white light for the indicated period. Wl +FR treatments were provided by supplementary FR LEDs to a R:FR ratio of 0.2.

#### Development of Transgenic Lines

*pDR5v2:GUS*-The *pGREENII0179:pDR5v2:GUS* plasmid was constructed by ligating a *pDR5v2* fragment cut from *pUC57:DR5v2* [19] with EcoRI and BamHI into *pGREENII0179* lacking a promoter. The *GUS* gene was amplified from a *pDR5:GUS* arabidopsis line containing a *GUS* reporter with primers introducing NotI and SacI restriction sites. The PCR fragment was subsequently cut with NotI and SacI and ligated into *pGREENII0179:pDR5v2* to create *pGREENII0179:pDR5v2:GUS*. This was then transformed into *E.coli* (strain-DH5α) and sequenced to confirm the correct insertion of *GUS*. The *pDR5v2:GUS* arabidopsis line was made transforming *pGREENII0179:pDR5v2:GUS* into *agrobacterium* (strain AGL1) containing the *pSOUP* plasmid. This was then transformed into Col-0 plants by floral dipping, according to an updated protocol [42]. 21 independent transformants were selected by antibiotic resistance and T2 lines screened for single insertions and GUS staining intensity. T3 homozygous lines were selected by kanamycin resistance and checked for signal strength, distribution and auxin responsiveness.

*pIAA19:LUC*-The *pIAA19:LUC* plasmid was constructed by amplifying the *IAA19* promoter from arabidopsis thaliana (Col ecotype), with the primers listed in the key resources table (see online). The promoter fragment was then cloned into *pENTR_D-TOPO* and mobilized into the LucTrap3 vector. This construct was then transformed into *agrobacterium* (strain AGL1) and used for transformation of Col-0 plants by floral dip.

*p35S:PIF4-HA* and *p35:PIF5-HA*-The *pCF402* and *pCF404* plasmids (described previously in [3]) were introduced into *agrobacterium* and then transformed into L*er* plants by floral dip. These lines are very similar to existing lines in the Col-0 background [3].

For oligonucleotides used in this study see Methods S1.

#### Gene Expression Studies

Seedlings were grown in the indicated conditions, before dissection and flash freezing in liquid nitrogen. RNA was extracted with an RNeasy Mini Kit (QIAGEN) with an on-column treatment with DNase (QIAGEN). cDNA was synthesized using a RevertAid First Strand cDNA Synthesis Kit (Thermo Fisher Scientific) with random hexamers. qPCR was performed with SYBR green ready mix (Bio-rad) in a ViiA 7 Real-Time PCR System (Thermo Fisher Scientific). Ct values for the gene of interest are expressed relative to that Ct value found for primers targeted to *AT1G13320*. For each experiment, 2 technical and a minimum of 3 biological repeats was performed. Primer efficiency tests were performed for each primer pair used.

For oligonucleotides used in this study see Methods S1.

#### Luciferase Assay

*pPIL1:LUC* and *pIAA19:LUC* seedlings were germinated in ½ MS media at 120-140 µmolm^-2^s^-1^ white light (16h:8h PP). On cotyledon expansion, seedlings were transferred to microtiter plates containing 170 µl solid ½ MS media (6% agar) supplemented with EtOH (mock), 75 mM NaCl (NaCl), or 2 µM ABA, per well. 30 µl of a luciferin solution (8 µl of a 10 mg/ ml luciferin (Promega) stock in DMSO, diluted in 4 ml H2O) were added per well, and plates were sealed with a sealing film (Applied Biosystems). Two holes were then made with a fine needle per well for seedlings transpiration. Plants were moved to red light (30-35 µmol m^-2^ s^-1^-16h:8h PP) and luminescence was recorded by a Berthold LB960 station installed in a Percival growth chamber. Luminescence was recorded each hour, with a 2 second read per well. On day 2 at Zt4, R light was supplemented with FR.

#### ABA Extraction and Quantification

Plants were grown under the same conditions used for morphological studies. At Zt 4 on day 7, 20-50mg of tissue (approximately 50 seedlings) were harvested and flash frozen in liquid nitrogen. Samples were homogenized (Precellys 24, Bertin Technologies, Aix-en-Provence, France) and extracted in ethyl acetate with an internal standard of D6-ABA (C/D/N Isotopes Inc, Canada). Samples were dried on a vacuum concentrator (CentriVap Centrifugal Concentrator, Labconco, Kansas City, MO, USA) at room temperature. Residues were re-suspended in 0.25 ml of 70% methanol (v/v) and samples were analyzed by liquid chromatography-mass spectrometry (LC-MS) on a Varian 320 Triple Quad LC-MS/MS. ABA levels were quantified from the peak area of each sample compared with the internal standard and normalized by fresh weight.

#### Western Blots

Plants were germinated on ½ MS plates in white light (120-140 µmol m^-2^ s^-1^ −16:8h PP) for 3 days, before being transferred to plates with or without 75mM NaCl. Plates were then grown in R light (30-35 µmol m^-2^ s^-1^ −16:8h PP) for a further 2 days. On day 5, at Zt 3, half the plates were moved to R+FR light. Tissues were harvested at Zt 4. 10 seedlings were homogenized and proteins were extracted in 70 µl Cracking Buffer (125mM Tris pH 7.4, 2% SDS, 10% Glycerol, 6M Urea, 5% βME) and 15 µl of the extract was run on a 10% polyacrylamide gel. Blots were probed with anti-HA-HRP (Roche) or anti-GFP-HRP (Miltenyi Biotec) as indicated in the text. Membranes were developed with 50/50 mix of ‘pico’ and ‘femto’ chemiluminescence substrates (Thermo) on a ChemiDoc (Biorad).

#### GUS Staining

Tissues were harvested an immediately fixed in 90% acetone at −20°C. Tissues were washed twice under a vacuum in GUS wash solution (0.1M Phospho-PI pH 7.0, 10 mM EDTA, 2 mM K_3_Fe(CN)_6_) and then stained overnight at 37°C in GUS staining solution (0.1M Phospho-PI pH 7.0, 10 mM EDTA, 1 mM K_3_Fe(CN)_6_, 1mM K_4_Fe(CN)_6_ * 3H_2_O, 0.5 mg/ ml X-glucuronide). The GUS reaction was stopped by treatment with 3:1 ethanol: acetic acid mix for 1 hour at 37°C. Tissues were cleared with 70% ethanol over several days and then mounted in 10% chloral hydrate, 30% glycerol. Representative images were taken with a Zeiss Axioskop 2 with Infinity 3 camera attachment and slides were scanned with an Epson v300 scanner for the purposes of GUS quantification.

### Quantification and Statistical Analyses

#### Image quantification

For morphological studies, plants were laid out on 0.8% agar plates and scanned with an Epson v300 scanner. Length measurements were then made in ImageJ. Western blot band intensities were quantified by measuring pixel intensity in ImageJ. Total protein loading was assessed through Rubisco Large Subunit band intensity, as visualized through ponceau stinging. Total protein loading was then used to normalize chemiluminescence blot band intensity. GUS staining quantification was performed in the ImageJ-based program, Icy. Relative GUS levels in the top half of the hypocotyl were defined by the mean intensity in Ch-0 minus the mean intensity in Ch-1.

#### Data presentation

Data presented as box and whisker diagrams were created in Graphpad Prism, and compiled in Microsoft Powerpoint. Data sets where n ≥ 10 are represented with a Tukey representation. The upper and lower edges of the box represent interquartile range (IQR) and the middle bar represents the median value. values 1.5 x IQR higher than the 75^th^ percentile or 1.5x IQR lower than the 25^th^ percentile are marked as single point outliers. Whiskers are plotted as the maximum and minimum values, or as 1.5x IQR, depending on which is closer to the median. Data sets where n ≤ 10 are presented as box and whisker diagrams, with all points shown. Whiskers represent the maximum and minimum values.

#### Statistical Analyses

1-way and 2-way ANOVAs were performed in Graphpad Prism or with SPSS statistics. T-tests were performed in Microsoft Excel. Significant differences are defined as p > 0.05. In graphs that include letters, each letter represents a statistically significant mean.

### Data Availability

The data used in the preparation of this manuscript, including morphological measurements, average Ct values, image analyses, ABA quantifications and photon counts is available at the following site: https://zenodo.org/record/2592526#.XIk36yhKg2w

The images used for western blot quantification are available at the sites listed below: PIF4-HA and PIF5-HA: https://zenodo.org/record/1343501#.XH_f2ohKg2w

BES1-GFP: https://zenodo.org/record/1480822#.XH_fCohKg2w

The images used for GUS-staining quantification are available at the site listed below:

*DR5v2:GUS* in both Col and *pif4pif5* mutants: https://zenodo.org/record/2022607#.XH_f74hKg2w

## Supplemental Item Titles

Methods S1: Primers used in this study (Related to Figures 2, S2, 3, S3, 4 and S4).

---

**Figure S1.**
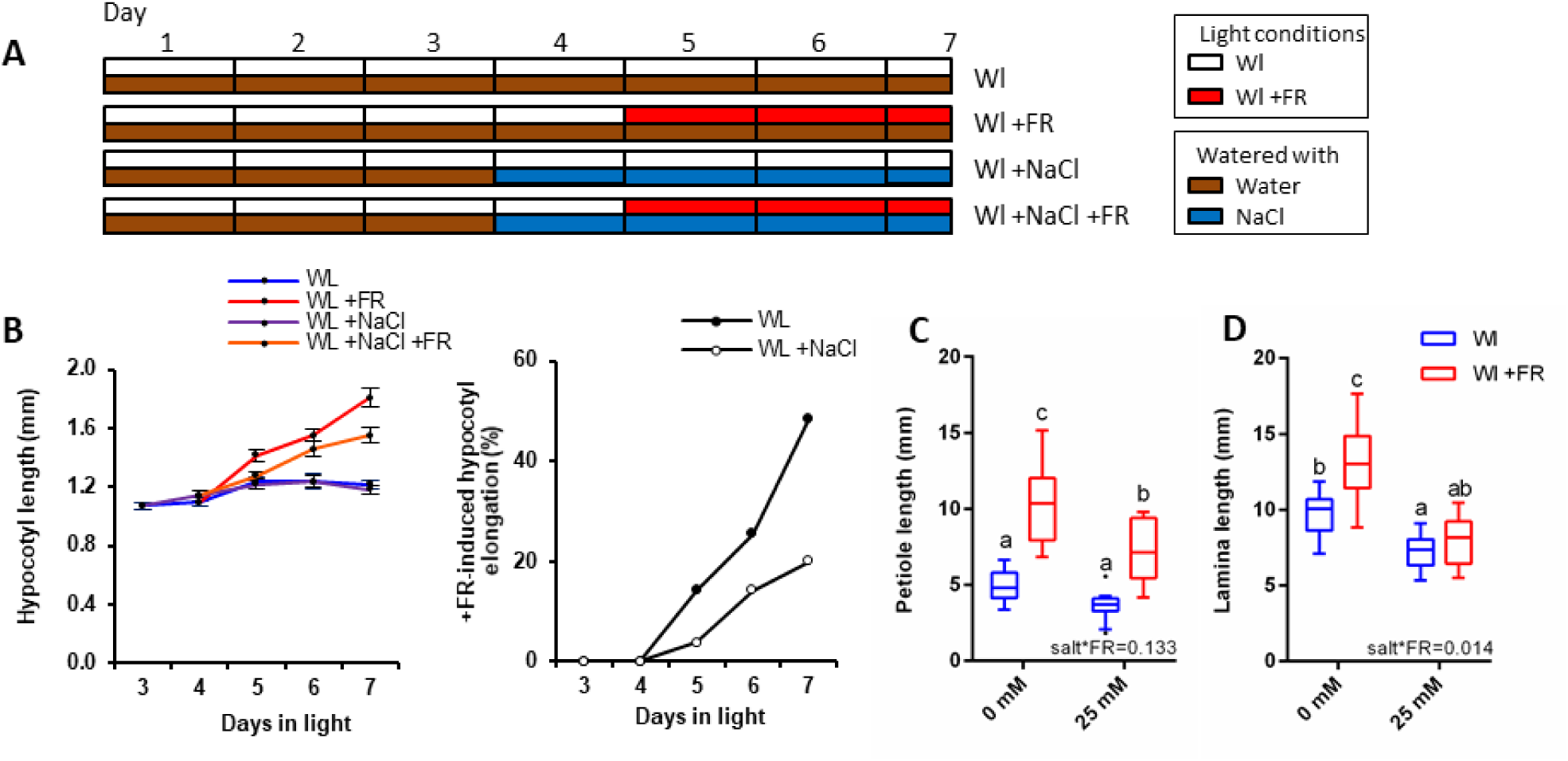
Soil salinity inhibits +FR-induced elongation; related to Figure 1. (**A**) Schematic representation of arabidopsis hypocotyl assay treatments. Plants were germinated on soil for 3 days before transfer to new pots that had been watered with water (Wl) or NaCI (Wl +NaCI). On day 4, half the plants were shifted into Wl +FR light (Wl +FR or Wl +NaCI +FR) and hypocotyls were measured on day 7. (**B**) Left: Hypocotyl length of Col seedlings seedlings grown germinated in Wl, before transfer to 25mM NaCI soil at day 3 and shifted into Wl +FR at day 4. Data represent mean (n>29) ± SE. Right: The same data expressed as the % +FR-induced elongation at each time point. (**C**) Petiole length or (**D**) lamina length of the 5^th^ leaf of the Col plants analysed in Figure IB (n>10).Box plots are visualised by the Tukey method. In all figures, different letters designate significantly different means by 2-way ANOVA + Tukey’s post-hoc test (p <0.05). Interaction P value is shown inset.*V

**Figure S2.**
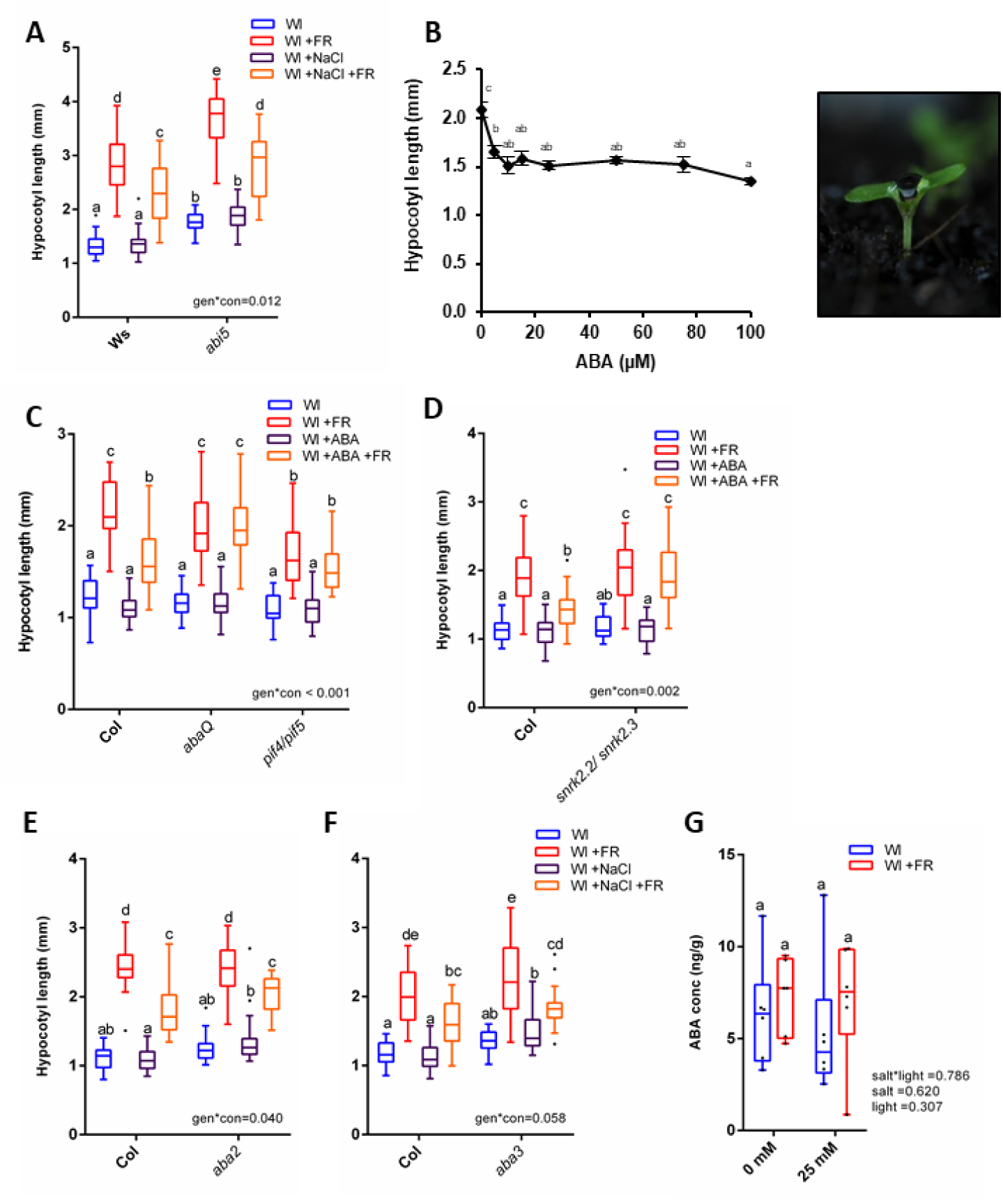
Salt inhibits +FR-induced hypocotyl elongation through ABA signalling, but this can still occur in several ABA mutants; related to Figure 2. (**A**) Hypocotyl length of 7-day old Ws and *abi5-l* seedlings germinated in Wl, transferred to ±25mM NaCI soil at day 3 and then shifted to WL ±FR at day 4 (n =30). (**B**) Left: Hypocotyl length of 7-day old Col seedlings germinated in Wl, transferred to new soil at day 3 and then shifted to WL +FR at day 4. The indicated concentration ABA was applied as 1 μl drops between the cotyledons on days 3-6 (n=24). Right: Demonstrative image of ABA treatments. (**C**) Hypocotyl length of 7-day old of Col, *abaQ* and *pif4-101/piß* or (**D**) Col and *snrk2.2/snrk2.3* seedlings germinated in Wl, transferred to new soil at day 3 and then shifted to WL ±FR at day 4.1 μl of 25 (iM ABA applied to each seedling on d 3-6 (n = 24). (**E**) Hypocotyl length of Col and *aba2-l* or (**F**) Col and *aba3-l* seedlings grown as in “A” (n >23). (**G**) ABA content of 7-day old Col seedlings grown as in “A” and harvested on day 7 at Zt 4 (n=6). Box plots are visualised by the Tukey method (n >10) or as max/min with every value shown (n < 10). Different letters designate significantly different means by 2-way ANOVA +Tukey’s post-hoc test (p <0.05). Interaction P value is shown inset.

**Figure S3.**
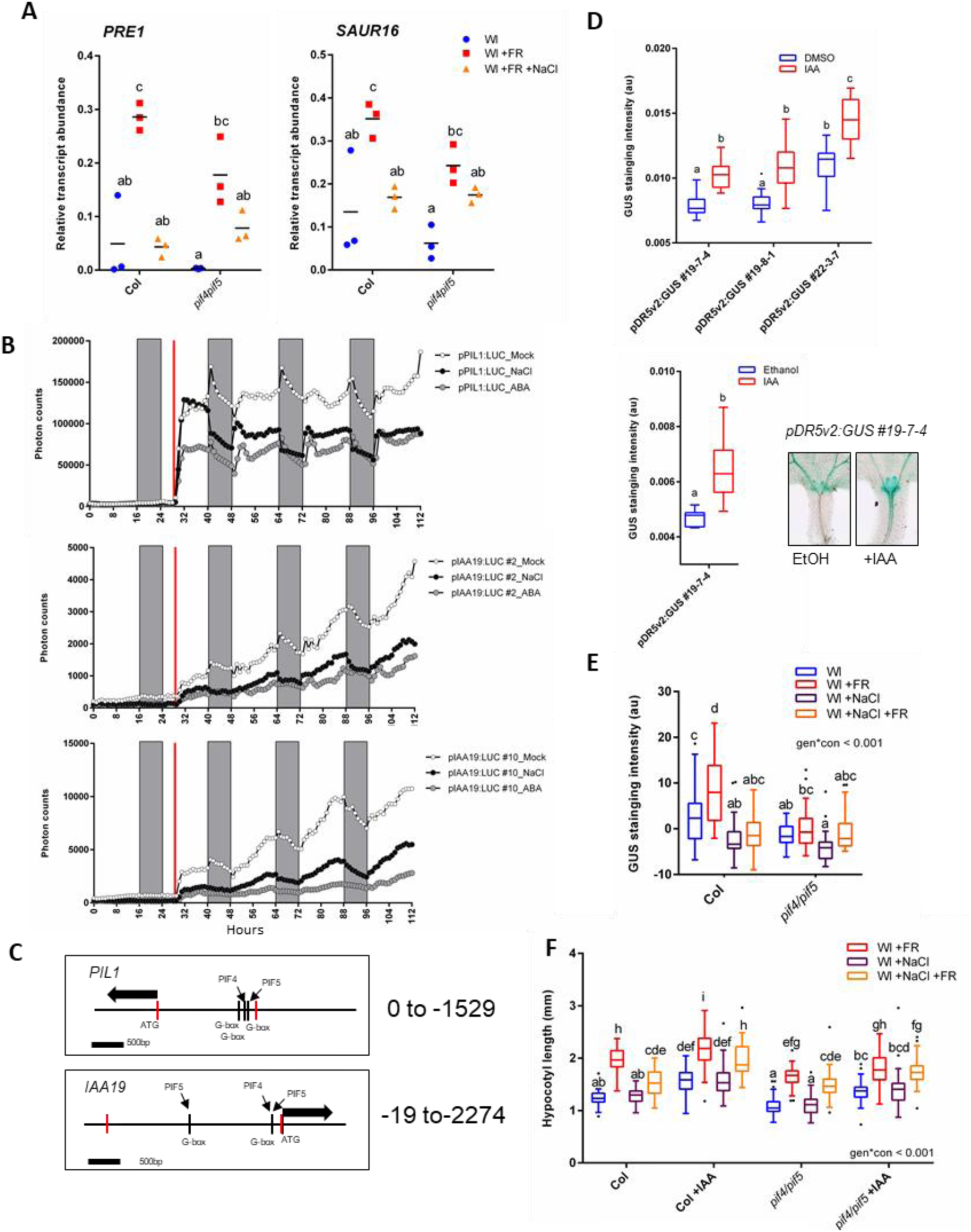
NaCI and ABA supress PIF action and auxin signalling; related to Figure 3. (**A**) Relative *PRE1* (left) and *SAUR16* (right) transcript abundance in the hypocotyls of 6 day old Col and *pif4pif5* mutants. Plants were germinated in Wl, transferred to ±25mM NaCI soil at day 3 and then shifted to WL ±FR at day 4. Tissues were harvested at Zt 4.5 on day 6 (n=3). (**B**) Seedlings expressing luciferase reporters under the control of the *PIL1* or *IAA19* promoter were germinated in Wl for 3 days, before transferto a 96 well plate containing half MS supplemented with luciferin, 75mM NaCI, 2 μM ABA and/or ethanol control. Luciferase activity was recorded for one day before +FR LEDs were switched on (red line) at Zt 4 on day 3. Luciferase activity was then recorded for further 3 days. Grey bars represent night (16:8 h photoperiod). (**C**) Schematic diagram of the genomic DNA used for the *pPIL1:LUC* and *plAA19:LUC* reporters. Promoter region cloned is represented between the red lines. ATG site, G-boxes and known PIF binding sites are highlighted (see refs [5,18]). (**D**) Top: Relative GUS staining in the hypocotyl of three independent 4-day old *pDR5v2:GUS* reporter lines grown on plates in the presence or absence of 5μM IAA (n >10). Bottom right: Relative GUS staining in the hypocotyl of *pDR5v2:GUS* #*19-7-4* reporter line grown on soil for 7 days in Wl and treated for 2 h with l00μM IAA at zt 2.5 (n=11) and Bottom left: representative GUS-stained seedlings. (**E**) Relative GUS staining over whole hypocotyls from plants expressing *pDR5v2*.-GUS in the Col or *pif4-101/pif5* mutant background, as related to figure 3.B (n > 29). (**F**) Hypocotyl length of 7 day-old Col and *pif4-101/pif5* mutants germinated in Wl, transferred to ±25mM NaCI soil at day 3 and then shifted to WL±FR at day 4. Plants were sprayed with 50 μM IAA or ethanol control on days 4-6 (n > 36).

**Figure S4.**
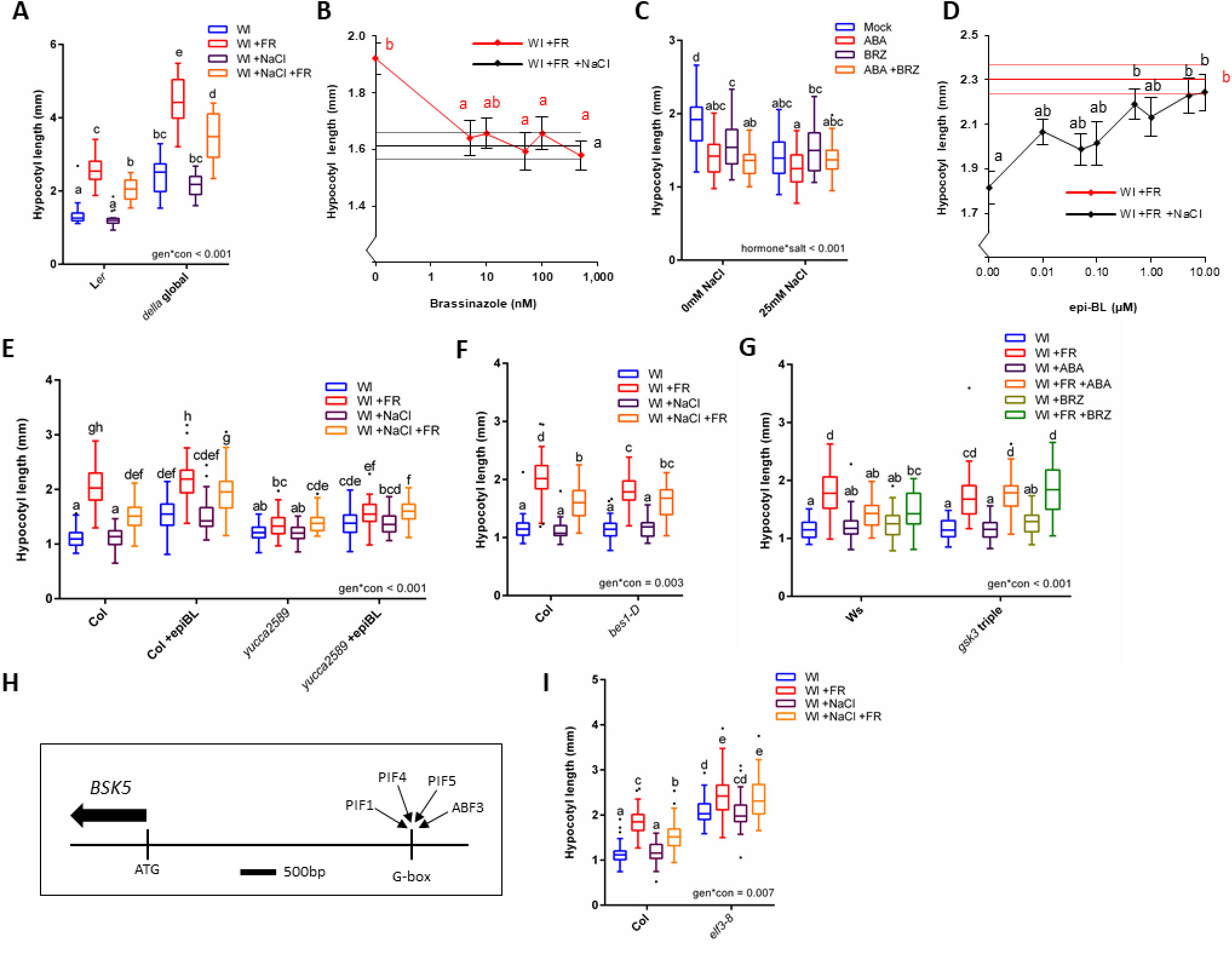
The inhibition of +FR-induced hypocotyl elongation by ABA and NaCI is mediated through a reduction in BR signalling; related to Figure 4. (**A**) Hypocotyl length of 7-day old L*er* and *della global* (pentuple) mutant germinated in Wl, transferred to ±25mM NaCI soil at day 3 and then shifted to WL ±FR at day 4 (n >17). (**B**) Hypocotyl length of 7-day old Col seedlings germinated in Wl, transferred to +25mM NaCI soil at day 3 and then shifted to WL+FR at day 4. 1 pi of the indicated concentration brassinazole was applied between the cotyledons on days 3-6. Data represent mean (n>52) ± SE. The black horizontal lines indicate the mean hypocotyl length (± SE) of control plants transferred to 25mM Nad soil and treated with mock applications; demonstrating that brassinazole inhibits hypocotyl elongation to the same extent as salt. (**C**) Hypocotyl length of 7-day old Col seedlings germinated in Wl, transferred to ±25mM NaCI soil at day 3 and then shifted to WL +FR at day 4. On days 3-6,1 μ l ethanol control, 25 μ M ABA, 50nM brassinazole (BRZ) or a combination of both was applied (n > 32). (**D**) Hypocotyl length of 7-day old Col seedlings germinated in Wl, transferred to ±25mM NaCI soil at day 3 and then shifted to WL +FR at day 4. The indicated concentration epi-brassinolide was sprayed onto the plants on days 3*6. Data represent mean (n>32) ± SE. The red horizontal lines indicate the mean hypocotyl length (± SE) of control plants transferred to non-Nad soil and treated with mock applications; demonstrating that epi-brassinolide promotes +FR-induced hypocotyl elongation to same extent as in non-salt treated plants. (**E**) Hypocotyl length of 7 day-old Col and *yucca2589* mutants germinated in Wl, transferred to +25mM NaCI soil at day 3 and then shifted to WL ±FR at day 4. Plants were treated with 10 pM epi-BL or ethanol control on days 3-6 (n > 45). (**F**) Hypocotyl length of 7-day old Col and *besl-D* mutants germinated in Wl, transferred to ±25mM NaCI soil at day 3 and then shifted to WL ±FR at day 4 (n = 36). (**G**) Hypocotyl length of 7-day old Ws and *gsk3-triple* seedlings grown as in “E” with 1 pi applications of 25 pM ABA or 50nM BRZ on d 3-6 (n > 25). (**H**) A schematic diagram of the BSK5 promoter, demonstrating G-boxes and PIF and ABF binding peaks identified in reference 12 and 40, respectively. (**I**) Hypocotyl length of 7-day old Col and *elf3-8* mutants grown as in “E” (n > 37). Box plots are visualised by the Tukey method. In all figures, different letters designate significantly different means by 2-way ANOVA +Tukey’s post-hoc test (p <0.05).

