## Supplementary material for "Soil salinity limits plant shade-avoidance": Key resources table

| **Reagent or Resource** | **Source** | **Identifier** |
| --- | --- | --- |
| Antibodies |  |  |
| High Affinity Anti-HA-HRP | Roche | 3F10 |
| Anti-GFP-HRP | Miltenyi Biotec | 130-091-833 |
| Bacterial strains |  |  |
| E.coli (DH5α) | n/a | n/a |
| A. tumefaciens (AGL1) | n/a | n/a |
| Chemicals, peptides and recombinant proteins |  |  |
| Murashige &Skoog (MS) medium | Duchefa Biochemie | M0222.0050 |
| Epibrassinolide | Santa-Cruz Biotechnology | sc-211419 |
| Abscisic Acid | Sigma Aldrich | 862169 |
| Brassinazole | Sigma Aldrich | SML1406 |
| Luciferin | Promega | E1601 |
| X-glucuronide | Sigma Aldrich | B5285-10MG |
| SuperSignal™ West Femto | Thermo Fisher Scientific | 34095 |
| SuperSignal™ West Pico PLUS | Thermo Fisher Scientific | 34580 |
| FastDigest SacI | Thermo Fisher Scientific | FD1133 |
| FastDigest NotI | Thermo Fisher Scientific | FD0593 |
| FastDigest EcoRI | Thermo Fisher Scientific | FD0274 |
| FastDigest BamHI | Thermo Fisher Scientific | FD0054 |
| FastDigest Green Buffer (10X) | Thermo Fisher Scientific | B72 |
| Critical Commercial Assays |  |  |
| RNeasy Mini Kit | QIAGEN | 74106 |
| RNase-Free DNase Set (50) | QIAGEN | 79254 |
| QIAprep Spin Miniprep Kit | QIAGEN | 27106 |
| Nucleospin Gel and PCR Clean-up | Macherey-Nagel | 740609 |
| RevertAid First Strand cDNA Synthesis Kit | Thermo Fisher Scientific | K1621 |
| SsoAdvanced™ Universal SYBR® Green Supermix | Bio-Rad | 1725271 |
| Deposited Data |  |  |
| PIF4/5-HA western blots | https://zenodo.org/record/1343501#.XH_f2ohKg2w | n/a |
| BES1-GFP western blots | https://zenodo.org/record/1480822#.XH_fCohKg2w | n/a |
| GUS staining images | https://zenodo.org/record/2022607#.XH_f74hKg2w | n/a |
| Raw data for figure generation | https://zenodo.org/record/2592526#.XIk36yhKg2w | n/a |
| Experimental organisms |  |  |
| Col-0 | n/a | n/a |
| Ws | n/a | n/a |
| L*er* | n/a | n/a |
| *pif4-101* | 3 | Garlic_114_G06 |
| *pif5 (pil6-1)* | 43 | SALK_087012 |
| *pif7-1* | 44 | SALK_044061 |
| *pif4-101/pif5* (aka *pil6-1*) | 3 | Garlic_114_G06/ SALK_087012 |
| *pif4-101/pif5* (aka *pil6-1*)/ *pif7-1* | 45 | Garlic_114_G06/ SALK_087012/ SALK_044061 |
| *abaQ* (*pyr1-1*/*pyl1-1*/*pyl2-1*/*pyl4-1*) | 14 | Point/ Salk_054640/ CSHL_GT2864_1/ Sail_517_C08 |
| *snrk2.2*/*snrk2.3* | 15 | GABI_807G04/ Salk_107315 |
| *arebQ (areb1/ areb2/ abf3/ abf1-1*) | 13 | SALK_002984/ SALK_069523/ SALK_096965/SALK_132819 |
| *abi5-1* | 46 | n/a |
| *bes1-D* | 27 | n/a |
| *gsk3-triple* (*bin2-3bil1-1bil2-1*) | 47 | FLAG_593C09/Wisonsin KO /Wisonsin KO |
| *bsk5* | NASC (characterised in 34) | SALK_051739C |
| *aba2-1* | 48 | n/a |
| *aba3-1* | 48 | n/a |
| *della global* (*gai-t6*/ *rga-t2*/ *rgl1-1*/ *rgl2-1*/ *rgl3-4*) | 49 | n/a |
| *pPIL1:LUC* | 17 | n/a |
| *pIAA19:LUC* | New line | n/a |
| *pDR5V2:GUS* | New line | n/a |
| *pDR5V2:GUS/ pif4-101/ pif5 (pil6-1)* | New line | n/a /Garlic_114_G06/ SALK_087012 |
| *p35S:PIF4-HA* | Franklin Lab, Bristol, UK | n/a |
| *p35S:PIF5-HA* | Franklin Lab, Bristol, UK | n/a |
| *p35S:BES1-GFP* | 28 | n/a |
| Tomato- S.lycopersicum var. “Moneymaker” | Intratuin, NL | EAN# 8717263349600 |
| Tobacco- N. benthamiana | n/a | n/a |
| Recombinant DNA |  |  |
| *pENTR_D-TOPO* | Thermo Fisher | K2400-20 |
| *LucTrap3* | n/a | GenBank: AY968054.1 |
| *pGREENII0179* | n/a | n/a |
| *pUC57:DR5v2* | Dolf Weijers (WUR) | n/a |
| Software and Algorithms |  |  |
| Microsoft Excel 2010 | Microsoft | n/a |
| Microsoft Powerpoint 2010 | Microsoft | n/a |
| Microsoft Word 2010 | Microsoft | n/a |
| SPSS Statistics 24 | IBM | n/a |
| Snapgene | GSL Biotech LLC | n/a |
| ImageJ | NIH | n/a |
| ICY | Quantitative Image Analysis Unit, Institut Pasteur | n/a |
| GIMP2 | The GIMP Development Team | n/a |
| ViiA™ 7 Software | Thermo Fisher Scientific | n/a |
| Prism 6 | Graphpad | n/a |
