## Supplementary material for "Soil salinity limits plant shade-avoidance": Oligonucleotides used in this study

### Slide 1
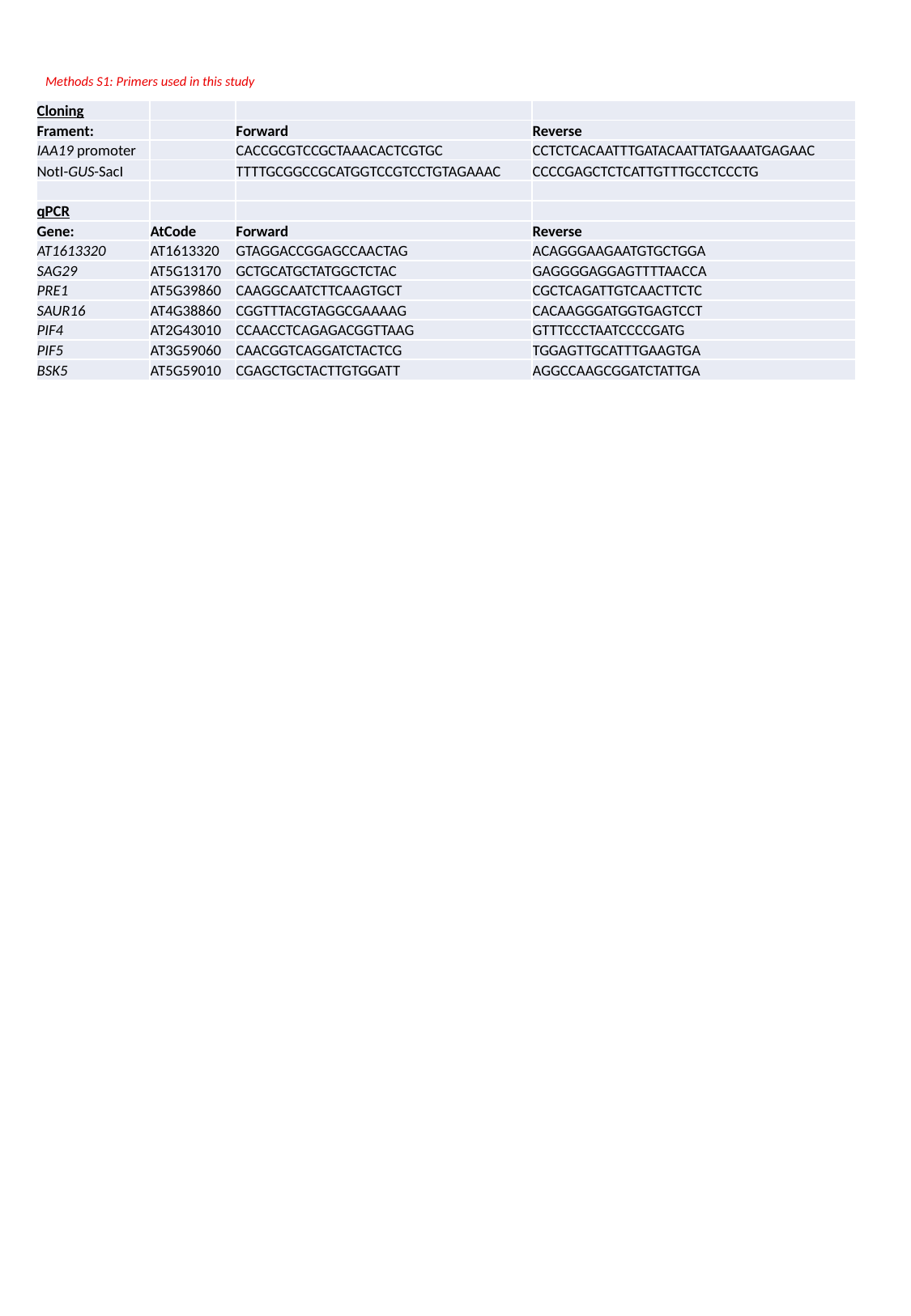

Methods S1: Primers used in this study
| Cloning | | | |
| --- | --- | --- | --- |
| Frament: | | Forward | Reverse |
| IAA19 promoter | | CACCGCGTCCGCTAAACACTCGTGC | CCTCTCACAATTTGATACAATTATGAAATGAGAAC |
| NotI-GUS-SacI | | TTTTGCGGCCGCATGGTCCGTCCTGTAGAAAC | CCCCGAGCTCTCATTGTTTGCCTCCCTG |
| qPCR | | | |
| Gene: | AtCode | Forward | Reverse |
| AT1613320 | AT1613320 | GTAGGACCGGAGCCAACTAG | ACAGGGAAGAATGTGCTGGA |
| SAG29 | AT5G13170 | GCTGCATGCTATGGCTCTAC | GAGGGGAGGAGTTTTAACCA |
| PRE1 | AT5G39860 | CAAGGCAATCTTCAAGTGCT | CGCTCAGATTGTCAACTTCTC |
| SAUR16 | AT4G38860 | CGGTTTACGTAGGCGAAAAG | CACAAGGGATGGTGAGTCCT |
| PIF4 | AT2G43010 | CCAACCTCAGAGACGGTTAAG | GTTTCCCTAATCCCCGATG |
| PIF5 | AT3G59060 | CAACGGTCAGGATCTACTCG | TGGAGTTGCATTTGAAGTGA |
| BSK5 | AT5G59010 | CGAGCTGCTACTTGTGGATT | AGGCCAAGCGGATCTATTGA |
